# Beetles provide directed dispersal of viable spores of a keystone wood decay fungus

**DOI:** 10.1101/2022.03.14.484227

**Authors:** Lisa Fagerli Lunde, Lynne Boddy, Anne Sverdrup-Thygeson, Rannveig M. Jacobsen, Håvard Kauserud, Tone Birkemoe

## Abstract

Wood decay fungi are considered to be dispersed by wind, but dispersal by animals may also be important, and more so in managed forests where dead wood is scarce. We investigated whether beetles could disperse spores of the keystone species *Fomitopsis pinicola*. Beetles were collected on sporocarps and newly felled spruce logs, a favourable habitat for spore deposition. Viable spores (and successful germination) of *F. pinicola* were detected by dikaryotization of monokaryotic bait mycelium from beetle samples. Viable spores were on the exoskeleton and in the faeces of all beetles collected from sporulating sporocarps. On fresh spruce logs, nine beetle species transported viable spores, of which several bore into the bark. Our results demonstrate that beetles can provide directed dispersal of wood decay fungi. Potentially, it could contribute to a higher persistence of some species in fragmented forests where spore deposition by wind on dead wood is less likely.

## 1. Introduction

In the boreal forests of Fennoscandia, many species of wood decay fungi are threatened with extinction due to intensive forest management, and subsequent habitat degradation and fragmentation (Kotiranta *et al*. 2019; SLU 2020; Brandrud *et al*. 2021). The declines have been linked to dispersal limitation, i.e. a mismatch between a species’ dispersal strategies and the altered landscape patterns in contemporary managed forests (Kallio 1970; Edman, Kruys & Jonsson 2004; Norros *et al*. 2012; Moor *et al*. 2021). Dispersal, defined as the movement by individuals or of propagules from their place of origin, is important for: (1) expanding or maintaining geographical ranges of populations, for instance by continuously establishing at new habitats (Hubbell 2001; Vellend 2010); and (2) maintaining or increasing genetic variability within the population through gene flow (Wright 1969; Ronce 2007). Dispersal of plants and fungi involves three separate stages, that can be likened to aviation (Ingold 1971): liberation (take-off), transport (flight), and deposition (landing). After that, albeit not defined as dispersal *per se*, the propagule needs to establish at the new habitat. While movement is relatively easy to observe for large animals, studying dispersal of microscopic propagules, such as fungal spores, is very challenging.

The most important wood decayers belong to the phylum Basidiomycota (Fungi), which inhabit the wood as mycelium. Spores are liberated from the sporocarp (fruit body) by the formation of a surface tension catapult (Buller 1933; Pringle *et al*. 2005). Then, spores of most species are transported by wind, although dispersal by animals or water also occurs (Ingold 1971; Birkemoe *et al*. 2018). Dead wood is patchily distributed, and the chances of landing on a suitable habitat when carried by wind currents are small because most dispersal propagules fall within a few meters of the source (Wenny 2001; Galante, Horton & Swaney 2011; Norros 2013). Wood decay fungi therefore produce an enormous number of spores, some liberating billions of spores per day (Ingold 1971). However, long-distance dispersal is considered to occur only rarely (Galante, Horton & Swaney 2011; Norros 2013; Golan & Pringle 2017), partly because airborne spores are highly vulnerable to abiotic stress (Kallio 1970; Norros *et al*. 2015).

Directed dispersal takes place when propagules arrive more often at favourable habitats than would be expected from random dispersal. In seed dispersal systems, directed dispersal is almost exclusively via animals (Wenny 2001) because they move in predictable ways, for example by foraging or dwelling in habitats with specific characteristics (Aukema & Martínez del Rio 2002; Leal, Wirth & Tabarelli 2007). Indeed, plants whose seeds are dispersed by animals are less vulnerable to fragmentation than wind-dispersed plants (Montoya *et al*. 2008; Sutton & Morgan 2009; Saar *et al*. 2012), which illustrates the potential advantages of directed dispersal.

Many species of fungi have adapted to dispersal by by animals, for example several tree pathogens (Webber & Gibbs 1989; Viiri 2007) and truffles (Trappe & Claridge 2005; Vašutová *et al*. 2019). Animal-mediated spore dispersal is also common among some wood decay fungi like odour-emitting stinkhorns (Ingold 1971; Tuno 1998), and those within obligate mutualisms, including with bark beetles and woodwasps. Bark beetles possess mycangia for carrying their fungal symbionts (Klepzig & Six 2004), but may occassionally also transport other wood decay fungi, such as *Fomitopsis pinicola* (Castello, Shaw & Furniss 1976; Pettey & Shaw 1986; Lim *et al*. 2006; Persson *et al*. 2009). The female woodwasp *Sirex noctilio* carries the white rot *Amylostereum areolatum* in mycangia and, after targeting a suitable tree, inoculates it directly into the wood (Slippers, De Groot & Wingfield 2011). Thus, she effectively aids all stages of the fungus’ dispersal – liberation, transport and deposition.

Dispersal of wood decay fungi by non-mutualistic invertebrates may be underestimated (e.g. Talbot 1952; Persson *et al*. 2009). Based on DNA analyses, a high diversity of wood decay fungi was found on saproxylic beetles (Jacobsen *et al*. 2017; Seibold *et al*. 2019) and on the red-cockaded woodpecker (Jusino *et al*. 2016). While detection of DNA is far from evidence of dispersal, successful transfer of viable spores has been demonstrated, for example by true flies (Diptera) in *Ganoderma* spp. (Lim 1977; Nuss 1982; Tuno 1999). Theoretically, the most beneficial type of animal vector for an early-colonizing wood decay fungus would deposit spores on fresh dead wood, for instance on a dying or recently dead tree. Many saproxylic beetles, in particular bark beetles, seek fresh dead wood, bore through the bark and excavate galleries in the vascular cambium to lay their eggs, thus creating a potential entry point for fungi (Ehnström & Axelsson 2002). Even beetles that do not bore through the bark, frequently seek shelter in cavities on or under the outer bark, thereby potentially assisting fungal spores in by-passing the bark barrier (Dossa *et al*. 2018).

In this study, we investigate whether beetles disperse spores of wood decay fungi, without being involved in mutualistic symbiosis. As a model species, we use *Fomitopsis pinicola* (Sw.) P. Karst, a common and widespread brown rot fungus in Fennoscandia. In boreal forests, it is a keystone species that contributes greatly to decomposition (Mounce 1929; Gramss 2020), and is an important predecessor of the red-listed fungi *Pycnoporellus fulgens, Antrodiella citrinella* (Niemelä, Renvall & Penttilä 1995; Norberg *et al*. 2019) and beetle *Peltis grossa* (Weslien *et al*. 2011). It is often an early colonizer of spruce and its sporocarps are habitats to a range of other species (Komonen *et al*. 2004; Lunde *et al*. 2022; Birkemoe et al. in prep.). For successful dispersal, spores need to be liberated, transported and deposited at a favourable habitat, and then successfully germinate. Therefore, we ask: (1) Do beetles that visit sporulating sporocarps of *F. pinicola* carry viable spores? We hypothesize that spore viability is greater when carried on the beetle exoskeleton than in the faeces, due to destruction of spores during digestion. (2) Do beetles that visit newly felled spruce logs carry viable spores of *F. pinicola*? Answering these questions will increase our understanding of dispersal strategies and potential dispersal limitation of wood decay fungi in general, which is integral to their conservation in contemporary managed forests.

## 2. Materials and methods

### 2.1. Study area

The study was set in the Nordre Pollen nature reserve in Southeastern Norway from May to June 2020 and 2021. Collection of beetles (as described below) was carried out at a spruce-dominated forest stand around Pollefløyta (59°75’N 10°76’E, 38 m.a.s.l; ∼500 m radius). On 10 March 2021, one spruce tree (*Picea abies* (L.) H. Karst.) was felled in the nearby Ås municipality (59°66’N 10°79’E, 92 m.a.s.l). The tree was cut into 15 ∼1 m long logs (diameter 17.4 – 27.9 cm) and placed at Pollefløyta on 29 April 2021 for beetle collection, in order to test their spore-vectoring capacity.

### 2.2. Spore baiting technique

*Fomitopsis pinciola* can grow, similar to most filamentous basidiomycete fungi, both as monokaryons and dikaryons. Mating begins with dikaryotization – i.e. when two monokaryons of compatible mating types fuse to form a dikaryon – and it ends when haploid spores are produced on sporocarps.

Spore dispersal is difficult to study in many wood decay fungi, such as *Fomitopsis pinicola*, because both mycelia and spores are hard to distinguish morphologically from related species. Furthermore, although plating onto artificial culture media can be used to demonstrate spore viability by germination and growth of mycelium, contamination from bacteria or other fungi is a common hindrance. To circumvent these issues, we used monokaryotic cultures of *F. pinicola* to trap spores from the exoskeleton and faeces of the collected beetles (Adams *et al*. 1984), an approach that has been previously used successfully to study wind dispersal in wood decay fungi (e.g. Williams, Todd & Rayner 1984; Edman, Kruys & Jonsson 2004; Boddy, Crockatt & Ainsworth 2011; Norros *et al*. 2012). Essentially, monokaryons were grown on agar media, and beetle heads, elytra, guts or faeces were placed individually on separate cultures. If viable spores of the same fungus species are present on the beetle samples, they can germinate and fuse with the monokaryons to form a dikaryon, provided that they are mating-type compatible (which will be the case for spores from almost all other sporocarps of the same species). Dikaryotization is evidenced microscopically by the presence of clamp connections, easily visible structures that form on the outside of hyphae so that the second nucleus can migrate between adjacent hyphal cells.

### 2.3. Monokaryon isolation

To isolate a monokaryotic strain to use as bait mycelium in the following experiments, five living sporocarps of *Fomitopsis pinicola* were collected from the study area (24 February 2020) and incubated in a room with a temperature of 30°C and high humidity. After 1 day, glass slides were placed underneath the hymenium for 2 hours to collect spores. Sterile, deionized water was mixed with the spore prints and added onto 55 mm Petri dishes containing 2% malt agar solution in a dilution series. After 3 – 4 days, single germinated spores were removed onto fresh agar using a sterile inoculation loop and a stereomicroscope. Two strains, originating from different sporocarps, were isolated and identified as monokaryons by absence of clamp connections and successful mating with each other. The strain with the fastest growth was chosen to use as bait in the following experiments with beetles.

### 2.4. Collection of beetles

To determine whether beetles visiting *F. pinicola* sporocarps could carry viable spores on their bodies or in their faeces, eight species were collected between 4 May and 9 June 2020 (Table 2). The beetles were collected after sunset, between 9.30 p.m. and 1.30 a.m, in 2 mL Eppendorf tubes which were then placed in a cool container to anesthetize them. A spore print was obtained from every sporocarp with a beetle, by placing a piece of adhesive tape on the spot where the beetle had been collected. The spore print was used to verify that the sporocarp had been sporulating. To determine whether viable spores survived the digestive tract, three of the beetles species from sporocarps (above), were collected between 2 July and 9 July 2021. The collected beetles were immediately submerged in sterile physiological saline water with a drop of soap to break the surface tension.

To determine whether beetles could transport viable spores of *F. pinicola* to a suitable habitat for deposition, 15 spruce logs from a recently felled tree were placed in the study area on 29 April 2021. Part of the log was covered with black drainage board to act as a shelter for potential beetles. All beetles that were found on the logs were collected during eleven separate visits, mostly during the day, from 4 May to 11 June 2021. As in 2020, the beetles were collected in 2 mL Eppendorf tubes and kept cold.

### 2.5. Detection of viable spores on beetle exoskeleton, guts and faeces

To detect viable spores from the collected beetles, living beetles were placed in glass Petri dishes that had been dry-heat sterilized for 30 min at 185°C. The room was 25°C with 16 h daylight (cool white fluorescent tubes), and beetles in Petri dishes were shaded with paperboard and given sterile deionized water on the tip of a cotton bud. The beetles in Petri dishes were left for 36 hrs, then refrigerated at 4°C for 1 h to lower their metabolism. Elytra, head and faeces were obtained aseptically (flame and ethanol sterilization) in a laminar flow cabinet with the use of a stereomicroscope, a scalpel, spring scissors and fine forceps, as described below. All faecal pellets that had been dropped in the Petri dish during the 36 hrs were collected. Care was taken to distinguish faeces from regurgitation matter or exoskeleton debris, although no investigations were made to verify that this was done successfully. During each round of sample preparation, a monokaryotic culture, in a Petri dish with the lid removed, was placed in the laminar flow cabinet as a control to test for contamination from ambient airborne *F. pinicola* propagules.

Sampling of beetles from the different collections varied slightly: (1) beetles from sporocarps (2020) were immediately killed by decapitation and three samples (faeces, elytra and head) were taken per individual; (2) from spruce logs (2021), two samples (faeces and one elytron) were taken per beetle individual and then submerged in 70% ethanol immediately after detaching the elytron, for species identification; (3) from sporocarps (2021), two samples (mid- and hindgut) were taken per beetle individual. In (3), the beetle and guts were thoroughly surface-sterilized; first, the beetle was transfered from the physiological saline solution to 70% ethanol for 10 s, then rinsed with sterile water. Second, guts were removed aseptically, submerged in 70% ethanol for 10 s and rinsed with sterile water. Finally, the mid- and hindgut were separated and punctured.

Each sample (elytra, head, faeces or gut) was placed directly on a monokaryotic *F. pinicola* culture, and the Petri dish was sealed with parafilm and incubated at 25°C in the dark for 20 – 21 days. Samples of the elytra and head (1 and 2) were rinsed with sterile deionized water, to remove potential contaminants from faeces, and patted dry on precision wipes before placing onto cultures. After incubation, each sample was subcultured onto a 55 mm diameter Petri dish with a 2% malt agar and antibiotic-fungicide solution (500 mg L^-1^ streptomycin, 300 mg L^-1^ penicillin, 5 mg L^-1^ benomyl [a fungicidal to most non-basidiomycetes]). About half of the samples from sporocarps (2020) were contaminated, mostly by secondary fungi (moulds), but they were successfully excluded during subculturing. After a few days, there was enough mycelial growth for microscopic examination. Hyphae with clamp connections were identified as dikaryons, indicating that the beetle sample had carried at least one viable spore of *F. pinicola*.

### 2.6 Beetle identification, data compilation and analysis

Beetles from sporocarps (2020 and 2021) were identified to species or genus level on site or in the lab by the authors. Beetles from spruce logs (2021) were sent to an expert for species identification. Metadata on diet, saproxylicity, favoured wood decay stage and boring behaviour were compiled for each species from the literature (Ehnström & Axelsson 2002; Seibold *et al*. 2015; SLU 2022). All data processing were done in R v 4.1.2 (Team 2021) and figures generated with the ggplot2 package (Wickham, Chang & Wickham 2016). To see whether exoskeleton samples (head or elytra) had statistically higher proportions of viable spores (detected by dikaryotization of bait mycelium) than faecal samples from beetles from sporocarps (2020), we used the χ^2^ contingency test (Wilson 1927). For beetle samples from spruce logs (2021), we used Fisher’s exact probability test because of small sample sizes (Fisher 1934).

## 3. Results

### 3.1. Do beetles that visit sporulating sporocarps of *F. pinicola* carry viable spores?

From sporulating sporocarps of *Fomitopsis pinicola*, we collected 167 beetles belonging to eight species in 2020 (Table 1). All beetle species carried viable spores of *F. pinicola* in their faeces and on their exoskeleton. Spores of *F. pinicola* survived passage through the digestive tract in 83.1% of all faecal samples, as detected by dikaryotization of monokaryotic bait mycelium (Figure 1). There were significantly more viable spores in head (0.98) and elytra (0.97) samples than in faecal (0.83) samples; χ^2^ = 17.87, p = < 0.001, and χ^2^ = 15.29, p = < 0.001, respectively (Figure 1a). To verify that the aforementioned result for faecal samples was not mere contamination from the exoskeleton, we dissected three of the species found on sporocarps (Table 1). Viable spores were detected in the guts of all three species, even after thorough surface sterilization.

**Table 1.** The number of faecal, elytra, head and gut samples from beetles (in alphabetical order) with viable spores of *Fomitopsis pinicola* (samples with viable spores/total number of samples), collected from sporulating sporocarps of *F. pinicola* and newly felled spruce logs. Information on taxonomy, saproxylicity, diet and preferred decay stage can be found in Table 2. *Two elytra were sampled from sporocarp beetles and one elytron from spruce log beetles.

| Species | Collected | Faeces | Elytra* | Head | Guts | Individuals |
| --- | --- | --- | --- | --- | --- | --- |
| *Ampedus nigrinus* | Spruce log | - | 0/1 | - | - | 1 |
| *Anisotoma humeralis* | Sporocarp | 9/11 | 19/20 | 19/19 | 2/2 | 20 |
|  | Spruce log | - | 0/1 | - | - | 1 |
| *Anthribus nebulosus* | Spruce log | - | 0/1 | - | - | 1 |
| *Atomaria vespertina* | Spruce log | 0/1 | 0/2 | - | - | 2 |
| *Corticaria longicollis* | Spruce log | - | 0/1 | - | - | 1 |
| *Dryocoetes autographus* | Spruce log | 0/2 | 1/2 | - | - | 2 |
| *Epuraea marseuli* | Spruce log | 0/1 | 0/2 | - | - | 2 |
| *Epuraea* sp. | Sporocarp | 11/16 | 20/21 | 22/22 | - | 24 |
| *Hylecoetus (Elateroides) dermestoides* | Spruce log | - | 1/2 | - | - | 2 |
| *Hylobius excavatus (piceus)* | Spruce log | 0/4 | 5/5 | - | - | 5 |
| *Hylurgops palliatus* | Spruce log | 0/2 | 1/4 | - | - | 4 |
| *Ipidia binotata* | Sporocarp | 21/33 | 42/43 | 41/42 | - | 44 |
| *Lordithon lunulatus* | Sporocarp | 3/4 | 8/8 | 8/8 | - | 8 |
| *Peltis ferruginea* | Sporocarp | 31/31 | 25/26 | 28/29 | 3/7 | 34 |
| *Placusa incompleta* | Spruce log | - | 0/1 | - | - | 1 |
| *Platynus assimilis* | Spruce log | - | 0/1 | - | - | 1 |
| *Polydrusus mollis* | Spruce log | - | 1/1 | - | - | 1 |
| *Pterostichus melanarius* | Spruce log | 0/1 | 0/3 | - | - | 3 |
| *Pterostichus oblongopunctatus* | Spruce log | 1/4 | 0/5 | - | - | 5 |
| *Ptinus subpilosus* | Spruce log | - | 0/1 | - | - | 1 |
| *Rhizophagus dispar* | Sporocarp | 2/2 | 1/1 | 2/2 | - | 2 |
|  | Spruce log | 0/3 | 0/3 | - | - | 3 |
| *Sepedophilus littoreus* | Sporocarp | 1/1 | 2/2 | 0/1 | - | 2 |
| *Sepedophilus testaceus* | Spruce log | 0/1 | 0/1 | - | - | 2 |
| *Stenichnus collaris* | Spruce log | - | 0/1 | - | - | 1 |
| *Tetropium castaneum* | Spruce log | - | 4/4 | - | - | 4 |
| *Thanasimus formicarius* | Spruce log | 0/5 | 5/8 | - | - | 8 |
| *Thymalus limbatus* | Sporocarp | 30/32 | 30/30 | 31/31 | 2/4 | 35 |
| *Trypodendron lineatum* | Spruce log | - | 3/5 | - | - | 5 |

**Figure 1.**
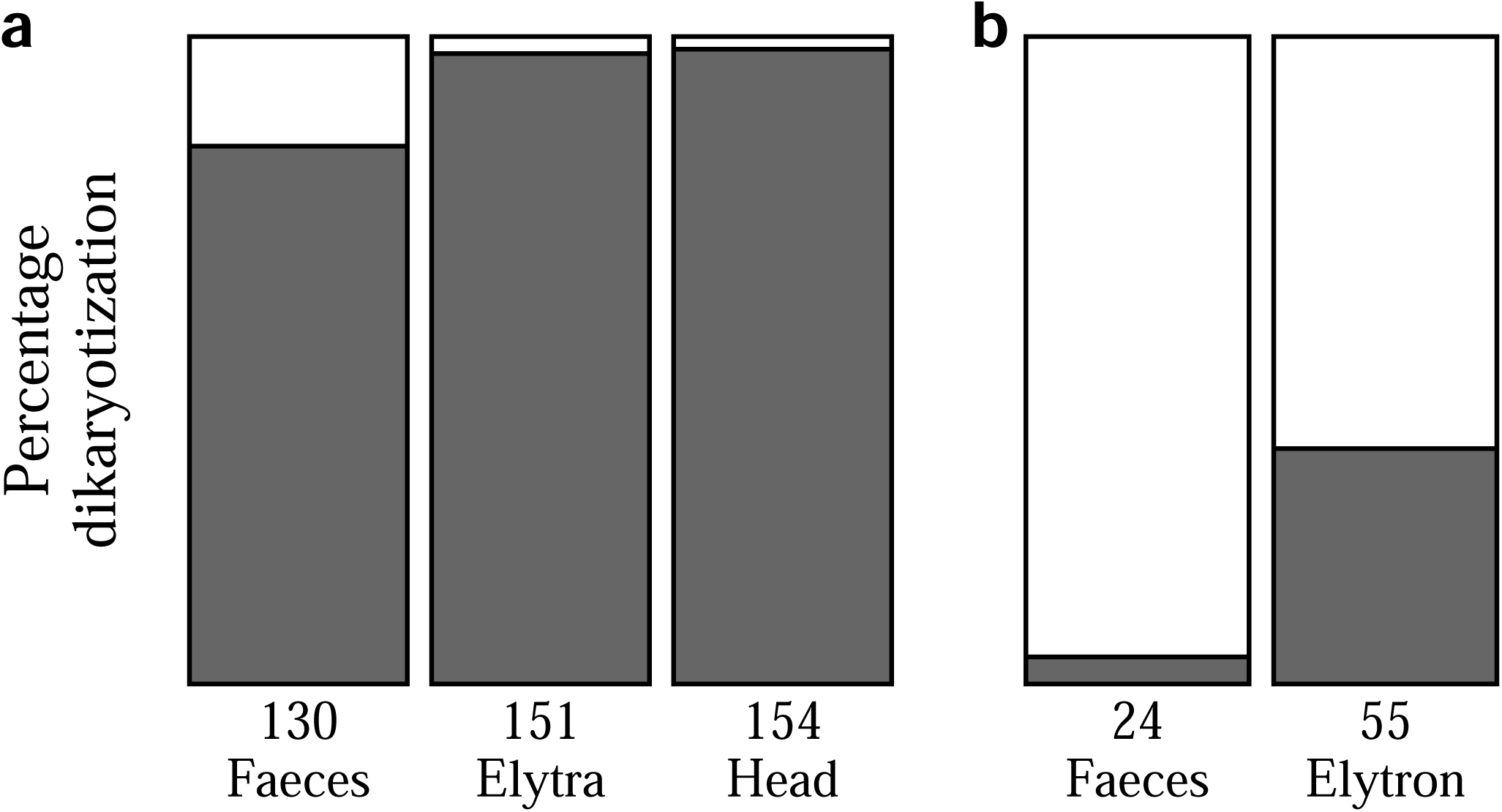
The proportion of viable spores (indicated by dikaryotization of bait mycelium, dark bars) of *Fomitopsis pinicola* (a) from beetle faecal, elytra and head samples collected from sporulating sporocarps (2020). (b) from beetle faecal and elytra samples collected from newly felled spruce logs (2021). Numbers below each bar represents the number of samples.

**Table 2.** Beetle species that were collected from sporulating sporocarps of *Fomitopsis pinicola* and newly felled spruce logs (Table 1), with information on taxonomy, diet, saproxylicity and preferred dead wood decay stage (1-5, from fresh to rotten), compiled from the literature (see Materials and Methods). Species in bold carried viable spores of *F. pinicola* on their exoskeletons or through their digestive tracts. 24 *Epuraea* sp. that were collected from sporocarps have been omitted because they were not identified to species. ^Bark beetles (Scolytinae).

| Species | Family | Diet | Saproxyllic | Preferred decay stage |
| --- | --- | --- | --- | --- |
| *Anthribus nebulosus* | Antribidae | Predator | Obligate |  |
| *Platynus assimilis* | Carabidae | Predator | No | --- |
| *Pterostichus melanarius* | Carabidae | Predator | No | --- |
| *Pterostichus oblongopunctatus* | **Carabidae** | Predator | **No** | **---** |
| *Tetropium castaneum* | **Cerambycidae** | **Xylophage** | **Obligate** | **2** |
| *Thanasimus formicarius* | **Cleridae** | **Predator** | **Obligate** | **2.25** |
| *Atomaria vespertina* | Cryptophagidae | Fungivore | No | --- |
| *Dryocoetes autographus*^ | **Curculionidae** | **Xylophage** | **Obligate** | **2** |
| *Hylobius excavatus* | **Curculionidae** | **Xylophage** | **Obligate** | **1** |
| *Hylurgops palliatus*^ | **Curculionidae** | **Xylophage** | **Obligate** | **2** |
| *Polydrusus mollis* | **Curculionidae** | **Herbivore** | **No** | **---** |
| *Trypodendron lineatum*^ | **Curculionidae** | **Fungivore** | **Obligate** | **2** |
| *Ampedus nigrinus* | Elateridae | Xylophage | Obligate | 3.67 |
| *Corticaria longicollis* | Lathridiidae | Fungivore | Facultative | 4.29 |
| *Anisotoma humeralis* | **Leiodidae** | **Fungivore** | **Obligate** | **4** |
| *Hylecoetus dermestoides* | **Lymexylidae** | **Xylophage** | **Obligate** | **2** |
| *Epuraea marseuli* | Nitidulidae | Predator | Obligate | 2.25 |
| *Ipidia binotata* | **Nitidulidae** | **Predator** | **Obligate** | **2.5** |
| *Ptinus subpilosus* | Ptinidae | Xylophage | Obligate |  |
| *Rhizophagus dispar* | **Rhizophagidae** | **Predator** | **Obligate** | **2.5** |
| *Lordithon lunulatus* | **Staphylinidae** | **Predator** | **Facultative** |  |
| *Placusa incompleta* | Staphylinidae | Predator | Obligate | 1.6 |
| *Sepedophilus littoreus* | **Staphylinidae** | **Fungivore** | **Facultative** |  |
| *Sepedophilus testaceus* | Staphylinidae | Fungivore | Facultative | 4.13 |
| *Stenichnus collaris* | Staphylinidae | Predator | No | --- |
| *Peltis ferruginea* | **Trogossitidae** | **Xylophage** | **Obligate** | **3.4** |
| *Thymalus limbatus* | **Trogossitidae** | **Xylophage** | **Obligate** | **3.25** |

### 3.2. Do beetles that visit newly felled spruce logs carry viable spores of *F. pinicola*?

We collected 55 beetles belonging to 22 species from newly felled spruce logs (Table 1). Eight species carried viable spores of *F. pinicola* on their elytra (36.4% of samples; Figure 1b) of which three were bark beetles (Table 2). The most common spore vectors were *Hylobius excavatus* (syn. *H. piceus*), *Thanasimus formicarius, Tetropium castaneum* and *Trypodendron lineatum* (Table 1). Only one faecal sample – from *Pterostichus oblongopunctatus* – had viable spores, which meant that elytra samples had a significantly higher odds ratio of carrying viable spores (one-tailed p = 0.0017). Beetle species carrying viable spores of *F. pinicola* on average preferred earlier decay stages of dead wood (mean 1.89, n = 7) than species that did not carry viable spores (mean 3.23, n = 6) (Figure 2).

**Figure 2.**
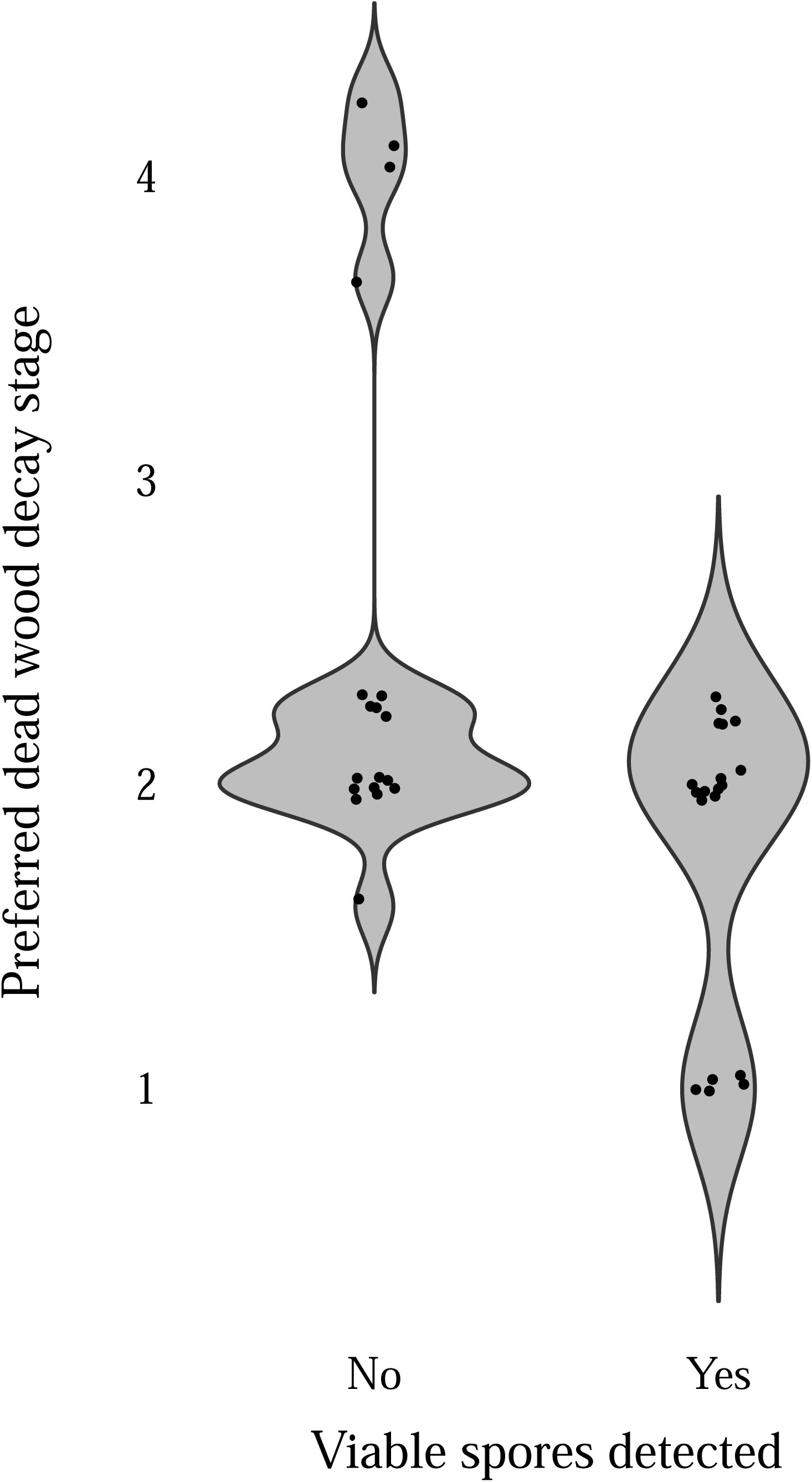
Violin plot displaying the preferred dead wood decay stage for 13 saproxylic beetle species and whether they had been carrying viable spores of *Fomitopsis pinicola* or not. Width of the violin is based on the relative kernal density estimate of the species at different decay stages.

## 4. Discussion

For an animal to be a successful vector of fungal spores, it needs to effectively aid in at least one of the stages of spore dispersal – liberation, transport and deposition. In addition, spores need to be deposited in favourable habitat and germinate. In this study, we have shown that beetles can disperse viable spores of *Fomitopsis pinicola*, even though these species are not involved in mutualistic symbiosis. We found viable spores both on the beetle exoskeleton, in the faeces and in the guts – thus, the spores even survived passage through the beetles’ digestive tracts. Specifically, all eight beetle species we collected on *F. pinicola* sporocarps carried spores that were viable at least 36 hours after collection. Among the 22 species collected from newly felled spruce logs, eight carried viable spores on the exoskeleton and one in the faeces. Our results demonstrate that beetles contribute to directed dispersal, with potential for successful deposition and establishment, of *F. pinicola* in new habitats.

Beetles that visit sporulating sporocarps physically remove spores, i.e. liberate, when they eat or walk on the spore-producing layer (hymenium). During our study period, we found many beetles visiting the sporocarps of *F. pinicola* at night, a well-known occurrence for saproxylic insects and wood decay fungi (e.g. Elton 1966), including *F. pinicola* (Hågvar 1999). While a diverse – and potentially specialized – community lives inside dead and living sporocarps (Jonsell & Nordlander 2004; Komonen *et al*. 2004; Lunde *et al*. 2022), both opportunistic and fungivorous insects visit to feed on the plentiful and nutritious spores. In this regard, it is tempting to draw analogies between insects visiting flowers to feed on nectar and pollen. In these interactions, plants attract insects to their flowers by volatiles to increase their chances of cross-pollination (Willmer 2011). Indeed, volatiles emitted from the sporocarps of some wood decay fungi act as attractants to visiting beetles (Jonsell & Nordlander 1995; Fäldt *et al*. 1999; Thakeow *et al*. 2008) and, in *Cryptoporus volvatus*, this kind of beetle attraction has been suggested to be involved in spore dispersal (discussed in Kües *et al*. 2018).

Beetles are active dispersers and wood decay fungi could potentially benefit from directed dispersal from beetles if they navigate to newly produced dead wood whilst carrying their spores. In the present study, nine beetle species transported viable spores of *F. pinicola* to newly felled spruce logs, which is likely to be a favourable habitat for spore deposition for the fungus. Saproxylic beetles collected from fresh dead wood logs have been found to carry wood decay fungi in the past (Jacobsen *et al*. 2017; Seibold *et al*. 2019). As these studies only detected DNA, however, the fungal propagules were not necessarily active or capable of establishing in the wood, which means that our study provides a valuable addition to our understanding of these interactions.

Three of the eight species that transported viable spores to newly felled spruce logs were bark beetles, which coincides with earlier studies that have isolated *F. pinicola* from two bark beetle genera (Castello, Shaw & Furniss 1976; Pettey & Shaw 1986; Lim *et al*. 2006; Persson *et al*. 2009). However, that these beetles carry spores, and not merely mycelial fragments from the decayed wood they emerge from as adults, was unexpected because bark beetles have not been observed to visit *F. pinicola* sporocarps in our study nor in others (Hågvar 1999, Birkemoe in prep.). For the bark beetles carrying viable spores to fresh spruce logs in the present study, spores were only found on the exoskeleton, and not in the faeces. Thus, these species might not feed on spores, but obtain them passively from the environment, for instance whilst dwelling in the vicinity of *F. pinicola* sporocarps. *Dryocoetes autographus* has been reported to be specifically attracted to wood colonized by *Fomitopsis rosea* (Johansson *et al*. 2006), while similar specificity has been shown for other beetles attracted to beech logs inoculated with different fungal species (Leather *et al*. 2014). As sporocarps of *F. pinicola* appear to be more abundant on logs that have been attacked by bark beetles (Pouska, Svoboda & Lepš 2013; Vogel *et al*. 2017), we could speculate that the beetles we collected from spruce logs might have obtained spores in aggregated patches of dead wood that are hotspots to both bark beetles and *F. pinicola*.

Spores of *F. pinicola* remained viable after passage through the digestive tracts of all the eight beetle species we collected from sporocarps. Internal dispersal of fungal spores has been demonstrated several times previously (e.g. Talbot 1952; Lim 1977; Colgan & Claridge 2002), including in beetles (Lilleskov & Bruns 2005; Drenkhan *et al*. 2013; Drenkhan, Kasanen & Vainio 2016). However, we found viable spores in the faeces of just one beetle from spruce logs – the non-saproxylic, predatory *Pterostichus oblongopunctatus* – that might have ingested spores indirecty through predation of spore-feeding invertebrates. Possibly, spores that are ingested by other beetle species at sporocarps are defaecated before the spore-feeders can transport them to a suitable habitat for deposition, or these beetles do not move to these habitats at all. In a different system, seeds of a parasitic plant (*Cytinus hypocistis*) were dispersed through beetle faeces, probably because they move to microsites that are favourable for seed deposition (de Vega *et al*. 2011). Such reliance on the navigation pattern of the vector, coupled with our results from spruce logs, suggest that internal spore dispersal by beetles, albeit possible, might be uncommon. However, our study is limited in spatial and temporal scale, and more research is needed to draw conclusions about the potential of internal spore dispersal by beetles in this system.

Even though beetles transport spores to suitable habitats, they are not necessarily deposited nor manage to establish there. The outer bark is an efficient physical barrier against invertebrates (Dossa *et al*. 2018), and it is usually intact in recently felled trees. Four of the species we detected transporting viable *F. pinicola* spores, however, including the three bark beetles and *Hylobius excavatus*, bore through the bark to lay their eggs in the cambium (Ehnström & Axelsson 2002). One of these species, *Hylurgops palliatus*, has been correlated with increased presence of *F. pinicola* sporocarps over 15 years (Weslien *et al*. 2011), but may reduce fungal diversity overall (Müller *et al*. 2002). Two other species that frequently carried viable spores, *Tetropium castaneum* and *Thanasimus formicarius*, prefer dead wood at early decay stages and lay eggs in cracks in the outer bark, but occasionally also go underneath the bark (Ye 1998; Ehnström 2007). Thus, these beetles could provide directed dispersal of *F. pinicola* spores not only to fresh logs, but past the bark barrier and into the wood, where the probability of spore establishment is higher. Based on our data, we cannot assert whether these interactions are overall beneficial to the fungus, but some bark beetles facilitate the establishment of wood decay fungi (Six 2013), including *F. pinicola* (Persson, Ihrmark & Stenlid 2011). Furthermore, several studies show that invertebrates essentially change the composition of wood decay communities during succession (e.g. Weslien *et al*. 2011; Jacobsen, Birkemoe & Sverdrup Thygeson 2015; Lunde in prep.). More research is needed to determine whether any of these effects on wood decay communities can be attributed to dispersal, and whether beetle dispersal affects fungal fitness overall.

To understand the measures that work for the conservation of wood decay fungi in boreal forests, it is crucial to know the abiotic and biotic factors they depend on, including their modes of dispersal. Moreover, wind dispersal has been considered to be *F. pinicola*’s predominant means of spread, but our results show that beetle dispersal could also be ubiquitous. This potential plasticity illustrates the importance of mapping different dispersal strategies for other species of wood decay fungi, as well. For instance, seed plants that are wind dispersed seem to be more vulnerable to habitat fragmentation and deforestation than plants whose seeds are dispersed by animals (e.g. Montoya *et al*. 2008). This might be true for wood decay fungi if animals provide a directed dispersal to the patchily distributed dead wood, but on the other hand, if a fungus depends on a few animal species for their dispersal, such dependency could make them more vulnerable to habitat fragmentation. Still, plants that depend on a few pollinators for cross-pollination have not been found to be more susceptible to habitat fragmentation (reviewed in Willmer 2011; Xiao *et al*. 2016). The results from seed dispersal or pollination systems do not necessarily translate to spore dispersal in wood decay fungi, for instance because most frugivorous animals are larger and move further than the typical spore feeder. Yet, investigating the relative importance of different dispersal strategies in wood decay fungi in general, and rare fungi in particular, is an essential step for understanding their potential responses to global threats, such as habitat fragmentation.

## Acknowlegdements

We would like to acknowledge Matthew Wainhouse for teaching fungal culturing; Mattias Edman and Sigmund Hågvar for discussions; Losby Bruk and Lars Juul for field permissions; Roar Økseter for spruce felling; Ross Wetherbee for metadata compilation; and Sindre Ligaard for beetle identification.

## Data availability

There is no additional data to deposit.

## References

Adams, T., Williams, E., Todd, N. & Rayner, A. (1984) A species-specific method of analysing populations of basidiospores. Transactions of the British Mycological Society, 82, 359–361.

Aukema, J.E. & Martínez del Rio, C. (2002) Where does a fruit-eating bird deposit mistletoe seeds? Seed deposition patterns and an experiment. Ecology, 83, 3489–3496.

Birkemoe, T., Jacobsen, R.M., Sverdrup-Thygeson, A. & Biedermann, P.H. (2018) Insect-fungus interactions in dead wood systems. Saproxylic insects, pp. 377–427. Springer.

Boddy, L., Crockatt, M.E. & Ainsworth, A.M. (2011) Ecology of Hericium cirrhatum, H. coralloides and H. erinaceus in the UK. Fungal ecology, 4, 163–173.

Brandrud, T., Bendiksen, E., Hofton, T., Jordal, J. & Nordén, J. (2021) Artsgruppeomtale sopper (Fungi). Norsk rødliste for arter 2021. Norwegian Biodiversity Information Centre (Artsdatabanken). https://www.artsdatabanken.no/Rodliste2021/Artsgruppene/Sopp Downloaded Jan 24 2022.

Buller, A. (1933) Researches on Fungi. Vol. V.. Longmans, Green and Co, London.

Castello, J.D., Shaw, C.G. & Furniss, M. (1976) Isolation of Cryptoporus volvatus and Fomes pinicola from Dendroctonus pseudotsugae. Phytopathology, 66, 1431–1434.

Colgan, W. & Claridge, A.W. (2002) Mycorrhizal effectiveness of Rhizopogon spores recovered from faecal pellets of small forest-dwelling mammals. Mycological Research, 106, 314–320.

de Vega, C., Arista, M., Ortiz, P.L., Herrera, C.M. & Talavera, S. (2011) Endozoochory by beetles: a novel seed dispersal mechanism. Annals of botany, 107, 629–637.

Dossa, G.G., Schaefer, D., Zhang, J.L., Tao, J.P., Cao, K.F., Corlett, R.T., Cunningham, A.B., Xu, J.C., Cornelissen, J.H. & Harrison, R.D. (2018) The cover uncovered: bark control over wood decomposition. Journal of Ecology, 106, 2147–2160.

Drenkhan, T., Kasanen, R. & Vainio, E.J. (2016) Phlebiopsis gigantea and associated viruses survive passing through the digestive tract of Hylobius abietis. Biocontrol Science and Technology, 26, 320–330.

Drenkhan, T., Sibul, I., Kasanen, R. & Vainio, E. (2013) Viruses of Heterobasidion parviporum persist within their fungal host during passage through the alimentary tract of Hylobius abietis. Forest Pathology, 43, 317–323.

Edman, M., Kruys, N. & Jonsson, B.G. (2004) Local dispersal sources strongly affect colonization patterns of wood-decaying fungi on spruce logs. Ecological Applications, 14, 893–901.

Ehnström, B. (2007) Skalbaggar: långhorningar. Coleoptera: Cerambycidae. Nationalnyckeln till Sveriges flora og fauna (The Encyclopedia of the Swedish Flora and Fauna). SLU Artdatabanken, Uppsala, Sweden.

Ehnström, B. & Axelsson, R. (2002) Insektsgnag i bark och ved. SLU Artdatabanken, Uppsala, Sweden.

Elton, C.S. (1966) Bracket Fungi and Toadstools. The Pattern of Animal Communities, pp. 306–318. Springer.

Fisher, R. (1934) Statistical Methods for Research Workers. Oliver and Boyd Ltd., Edinburgh.

Fäldt, J., Jonsell, M., Nordlander, G. & Borg-Karlson, A.-K. (1999) Volatiles of bracket fungi Fomitopsis pinicola and Fomes fomentarius and their functions as insect attractants. Journal of chemical ecology, 25, 567–590.

Galante, T.E., Horton, T.R. & Swaney, D.P. (2011) 95% of basidiospores fall within 1 m of the cap: a field-and modeling-based study. Mycologia, 103, 1175–1183.

Golan, J.J. & Pringle, A. (2017) Long-distance dispersal of fungi. Microbiology spectrum, 5, 5.4. 03.

Gramss, G. (2020) Aspects determining the dominance of Fomitopsis pinicola in the colonization of deadwood and the role of the pathogenicity factor oxalate. Forests, 11, 290.

Hubbell, S.P. (2001) The unified neutral theory of biodiversity and biogeography. Princeton University Press.

Hågvar, S. (1999) Saproxylic beetles visiting living sporocarps of Fomitopsis pinicola and Fomes fomentarius. Norwegian Journal of Entomology, 46, 25–32.

Ingold, C.T. (1971) Fungal spores. Their libération and dispersal. Oxford University Press, London.

Jacobsen, R.M., Birkemoe, T. & Sverdrup-Thygeson, A. (2015) Priority effects of early successional insects influence late successional fungi in dead wood. Ecology and Evolution, 5, 4896–4905.

Jacobsen, R.M., Kauserud, H., Sverdrup-Thygeson, A., Bjorbækmo, M.M. & Birkemoe, T. (2017) Wood-inhabiting insects can function as targeted vectors for decomposer fungi. Fungal ecology, 29, 76–84.

Johansson, T., Olsson, J., Hjältén, J., Jonsson, B.G. & Ericson, L. (2006) Beetle attraction to sporocarps and wood infected with mycelia of decay fungi in old-growth spruce forests of northern Sweden. Forest Ecology and Management, 237, 335–341.

Jonsell, M. & Nordlander, G. (1995) Field attraction of Coleoptera to odours of the wood-decaying polypores Fomitopsis pinicola and Fomes fomentarius. Annales Zoologici Fennici, pp. 391–402. JSTOR.

Jonsell, M. & Nordlander, G. (2004) Host selection patterns in insects breeding in bracket fungi. Ecological Entomology, 29, 697–705.

Jusino, M.A., Lindner, D.L., Banik, M.T., Rose, K.R. & Walters, J.R. (2016) Experimental evidence of a symbiosis between red-cockaded woodpeckers and fungi. Proceedings of the Royal Society B: Biological Sciences, 283, 20160106.

Kallio, T. (1970) Aerial distribution of the root-rot fungus Fomes annosus (Fr.) Cooke in Finland.

Klepzig, K.D. & Six, D. (2004) Bark beetle-fungal symbiosis: context dependency in complex associations. Symbiosis.

Komonen, A., Jonsell, M., Økland, B., Sverdrup-Thygeson, A. & Thunes, K. (2004) Insect assemblage associated with the polypore Fomitopsis pinicola: a comparison across Fennoscandia. Entomologica Fennica, 15, 102–112-102–112.

Kotiranta, H., Junninen, K., Halme, P., Kytövuori, I., T., v.B., Niskanen, T. & K., L. (2019) Aphyllophoroid fungi. pp. 234-247. Ministry of the Environment and Finnish Environment Institute, Helsinki. In: Hyvärinen, E., Juslén, A., Kemppainen, E., Uddström, A., Liukko, U.-M. (Eds.). The 2019 Red List of Finnish Species. Downloaded 24 Jan 2022.

Kües, U., Khonsuntia, W., Subba, S. & Dörnte, B. (2018) Volatiles in communication of Agaricomycetes. Physiology and genetics, pp. 149–212. Springer.

Leal, I.R., Wirth, R. & Tabarelli, M. (2007) Seed dispersal by ants in the semi-arid Caatinga of north-east Brazil. Annals of botany, 99, 885–894.

Leather, S.R., Baumgart, E.A., Evans, H.F. & Quicke, D.L. (2014) Seeing the trees for the wood–beech (Fagus sylvatica) decay fungal volatiles influence the structure of saproxylic beetle communities. Insect Conservation and Diversity, 7, 314–326.

Lilleskov, E.A. & Bruns, T.D. (2005) Spore dispersal of a resupinate ectomycorrhizal fungus, Tomentella sublilacina, via soil food webs. Mycologia, 97, 762–769.

Lim, T. (1977) Production, germination and dispersal of basidiospores of Ganoderma pseudoferreum on Hevea. Journal of the rubber Research Institute of Malaysia, 25, 93–99.

Lim, Y.W., Kim, J.-J., Lu, M. & Breuil, C. (2006) Determining fungal diversity on Dendroctonus ponderosae and Ips pini affecting lodgepole pine using cultural and molecular methods.

Lunde, L.F., Birkemoe, T., Kauserud, H., Boddy, L., Jacobsen, R.M., Morgado, L., Sverdrup-Thygeson, A. & Maurice, S. (2022) DNA metabarcoding reveals host-specific communities of arthropods residing in fungal fruit bodies. Proceedings of the Royal Society B, 289, 20212622.

Montoya, D., Zavala, M.A., Rodríguez, M.A. & Purves, D.W. (2008) Animal versus wind dispersal and the robustness of tree species to deforestation. Science, 320, 1502–1504.

Moor, H., Nordén, J., Penttilä, R., Siitonen, J. & Snäll, T. (2021) Long-term effects of colonization–extinction dynamics of generalist versus specialist wood-decaying fungi. Journal of Ecology, 109, 491–503.

Mounce, I. (1929) II. The biology of Fomes pinicola (Sw.) Cooke. Dominion of Canada Department of Agriculture, 111, 80.

Müller, M.M., Varama, M., Heinonen, J. & Hallaksela, A.-M. (2002) Influence of insects on the diversity of fungi in decaying spruce wood in managed and natural forests. Forest Ecology and Management, 166, 165–181.

Niemelä, T., Renvall, P. & Penttilä, R. (1995) Interactions of fungi at late stages of wood decomposition. Annales Botanici Fennici, pp. 141–152. JSTOR.

Norberg, A., Halme, P., Kotiaho, J.S., Toivanen, T. & Ovaskainen, O. (2019) Experimentally induced community assembly of polypores reveals the importance of both environmental filtering and assembly history. Fungal ecology, 41, 137–146.

Norros, V. (2013) Measuring and modelling airborne dispersal in wood decay fungi.

Norros, V., Karhu, E., Nordén, J., Vähätalo, A.V. & Ovaskainen, O. (2015) Spore sensitivity to sunlight and freezing can restrict dispersal in wood-decay fungi. Ecology and Evolution, 5, 3312–3326.

Norros, V., Penttilä, R., Suominen, M. & Ovaskainen, O. (2012) Dispersal may limit the occurrence of specialist wood decay fungi already at small spatial scales. Oikos, 121, 961–974.

Nuss, I. (1982) Die Bedeutung der proterosporen: Schlußfolgerungen aus untersuchungen an Ganoderma (Basidiomycetes). Plant systematics and evolution, 141, 53–79.

Persson, Y., Ihrmark, K. & Stenlid, J. (2011) Do bark beetles facilitate the establishment of rot fungi in Norway spruce? Fungal ecology, 4, 262–269.

Persson, Y., Vasaitis, R., Långström, B., Öhrn, P., Ihrmark, K. & Stenlid, J. (2009) Fungi vectored by the bark beetle Ips typographus following hibernation under the bark of standing trees and in the forest litter. Microbial ecology, 58, 651–659.

Pettey, T.M. & Shaw, C.G. (1986) Isolation of Fomitopsis pinicola from in-flight bark beetles (Coleoptera: Scolytidae). Canadian Journal of Botany, 64, 1507–1509.

Pouska, V., Svoboda, M. & Lepš, J. (2013) Co-occurrence patterns of wood-decaying fungi on Picea abies logs: does Fomitopsis pinicola influence the other species. Polish Journal of Ecology, 61, 119––133.

Pringle, A., Patek, S.N., Fischer, M., Stolze, J. & Money, N.P. (2005) The captured launch of a ballistospore. Mycologia, 97, 866–871.

Ronce, O. (2007) How does it feel to be like a rolling stone? Ten questions about dispersal evolution. Annu. Rev. Ecol. Evol. Syst., 38, 231–253.

Seibold, S., Brandl, R., Buse, J., Hothorn, T., Schmidl, J., Thorn, S. & Müller, J. (2015) Association of extinction risk of saproxylic beetles with ecological degradation of forests in Europe. Conservation Biology, 29, 382–390.

Seibold, S., Müller, J., Baldrian, P., Cadotte, M.W., Štursová, M., Biedermann, P.H., Krah, F.-S. & Bässler, C. (2019) Fungi associated with beetles dispersing from dead wood–Let’s take the beetle bus! Fungal ecology, 39, 100–108.

Six, D.L. (2013) The bark beetle holobiont: why microbes matter. Journal of chemical ecology, 39, 989–1002.

Slippers, B., De Groot, P. & Wingfield, M.J. (2011) The Sirex Woodwasp and its Fungal Symbiont:: Research and Management of a Worldwide Invasive Pest. Springer Science & Business Media.

SLU (2020) Rödlistade arter i Sverige 2020. Swedish Species Information Centre (SLU Artdatabanken), Uppsala.

SLU (2022) Artfakta (SLU, Artdatabanken). https://artfakta.se/artbestamning. Downloaded 10 Mar 2022.

Sutton, F.M. & Morgan, J.W. (2009) Functional traits and prior abundance explain native plant extirpation in a fragmented woodland landscape. Journal of Ecology, 97, 718–727.

Saar, L., Takkis, K., Pärtel, M. & Helm, A. (2012) Which plant traits predict species loss in calcareous grasslands with extinction debt? Diversity and Distributions, 18, 808–817.

Talbot, P. (1952) Dispersal of fungus spores by small animals inhabiting wood and bark. Transactions of the British Mycological Society, 35, 123–128.

Team, R.C. (2021) R: A language and environment for statistical computing.

Thakeow, P., Angeli, S., Weißbecker, B. & Schütz, S. (2008) Antennal and behavioral responses of Cis boleti to fungal odor of Trametes gibbosa. Chemical Senses, 33, 379–387.

Trappe, J.M. & Claridge, A.W. (2005) Hypogeous fungi: evolution of reproductive and dispersal strategies through interactions with animals and mycorrhizal plants. The Fungal Community: its organization and role in the ecosystem (eds J. Dighton & J.F. White), pp. 613–623. CRC Press.

Tuno, N. (1998) Spore dispersal of Dictyophora fungi (Phallaceae) by flies. Ecological Research, 13, 7–15.

Tuno, N. (1999) Insect feeding on spores of a bracket fungus, Elfvingia applanata (Pers.) Karst.(Ganodermataceae, Aphyllophorales). Ecological Research, 14, 97–103.

Vašutová, M., Mleczko, P., López-García, A., Maček, I., Boros, G., Ševčík, J., Fujii, S., Hackenberger, D., Tuf, I.H. & Hornung, E. (2019) Taxi drivers: the role of animals in transporting mycorrhizal fungi. Mycorrhiza, 29, 413–434.

Vellend, M. (2010) Conceptual synthesis in community ecology. The Quarterly review of biology, 85, 183–206.

Viiri, H. (2007) Fungi associated with Hylobius abietis and other weevils. Bark and wood boring insects in living trees in Europe, a synthesis, pp. 381–393. Springer.

Vogel, S., Alvarez, B., Bässler, C., Müller, J. & Thorn, S. (2017) The Red-belted Bracket (Fomitopsis pinicola) colonizes spruce trees early after bark beetle attack and persists. Fungal ecology, 27, 182–188.

Webber, J. & Gibbs, J. (1989) Insect dissemination of fungal pathogens of trees. Insect-fungus interactions, 160–175.

Wenny, D.G. (2001) Advantages of seed dispersal: a re-evaluation of directed dispersal. Evolutionary Ecology Research, 3, 37–50.

Weslien, J., Djupström, L.B., Schroeder, M. & Widenfalk, O. (2011) Long-term priority effects among insects and fungi colonizing decaying wood. Journal of Animal Ecology, 80, 1155–1162.

Wickham, H., Chang, W. & Wickham, M.H. (2016) Package ‘ggplot2’. Create Elegant Data Visualisations Using the Grammar of Graphics. Version, 2, 1–189.

Williams, E., Todd, N. & Rayner, A. (1984) Characterization of the spore rain of Coriolus versicolor and its ecological significance. Transactions of the British Mycological Society, 82, 323–326.

Willmer, P. (2011) Pollination and floral ecology. Princeton University Press.

Wilson, E.B. (1927) Probable inference, the law of succession, and statistical inference. Journal of the American Statistical Association, 22, 209–212.

Wright, S. (1969) Evolution and the genetics of populations: Vol. 2. The theory of gene frequencies.

Xiao, Y., Li, X., Cao, Y. & Dong, M. (2016) The diverse effects of habitat fragmentation on plant–pollinator interactions. Plant Ecology, 217, 857–868.

Ye, H. (1998) Life cycle of Thanasimus formicarius (Coleoptera: Cleridae) in Southern Norway. Insect Science, 5, 55–62.

